# LIN-10 can promote LET-23 EGFR signaling and trafficking independently of LIN-2 and LIN-7

**DOI:** 10.1101/2020.07.24.220293

**Authors:** Kimberley D. Gauthier, Christian E. Rocheleau

**Author notes:** Author to whom correspondence should be addressed Christian Rocheleau, Research Institute of the McGill University Health Centre, 1001 Décarie Blvd, Room E02.7242, Montreal, QC H4A 3J1. Abbreviations: CID, CASK-Interacting Domain; Muv, Multivulva; VPC, Vulva Precursor Cell; Vul, Vulvaless.

## Abstract

During *C. elegans* larval development, an inductive signal mediated by LET-23 EGFR (Epidermal Growth Factor Receptor) specifies three of six vulva precursor cells (VPCs) to adopt vulval cell fates. An evolutionarily conserved complex consisting of PDZ domain-containing scaffold proteins LIN-2 (CASK), LIN-7 (Lin7 or Veli), and LIN-10 (APBA1 or Mint1) (LIN-2/7/10) mediates basolateral LET-23 EGFR localization in the VPCs to permit signal transmission and development of the vulva. We recently found that the LIN-2/7/10 complex likely forms at Golgi ministacks or recycling endosomes; however, the mechanism through which the complex targets the receptor to the basolateral membrane remains unknown. Here we found that overexpression of LIN-10 or LIN-7 can compensate for loss of their complex components by promoting LET-23 EGFR signaling through previously unknown complex-independent and receptor-dependent pathways. In particular, LIN-10 can independently promote basolateral LET-23 EGFR localization, and its complex-independent function uniquely requires its PDZ domains that also regulate its localization to Golgi ministacks and recycling endosomes. These studies point to a novel complex-independent function for LIN-7 and LIN-10 that broadens our understanding of how this complex regulates targeted sorting of membrane proteins.

## Introduction

The hermaphroditic nematode *Caenorhabditis elegans* has served as an invaluable model in the discovery of the Epidermal Growth Factor Receptor (EGFR)/Ras/ERK signaling cascade, and in our general understanding of how cellular signaling and development are regulated. Development of the vulva, required for egg-laying and mating, is initiated by activation of the sole EGFR homolog, LET-23, in a set of progenitor epithelial cells called the vulva precursor cells (VPCs). The spatial organization of the signaling pathway within the VPCs is a critical dimension of signaling regulation: LET-23 EGFR must be present on the basolateral membrane of the VPCs to engage with the EGF-like ligand LIN-3, released from the overlying gonad, and to initiate vulval development (Aroian *et al.*, 1990; Hill and Sternberg, 1992; Simske *et al.*, 1996).

All six VPCs, named P3.p to P8.p, express LET-23 EGFR and can be induced into vulval cell fates; however, only the cell closest to the source of the ligand (P6.p) receives sufficient inductive signal to activate the downstream LET-60 Ras/MPK-1 ERK signaling cascade to specify the primary (1°) vulval cell fate (Sternberg and Horvitz, 1986; Aroian *et al.*, 1990; Han *et al.*, 1990). P6.p in turn signals to adjacent P5.p and P7.p cells through LIN-12 Notch signaling to induce them to adopt a secondary (2°) cell fate (Figure 1A), aided in part by graded inductive signaling through an alternate Ras/Ral GTPase pathway downstream of LET-23 EGFR (Greenwald *et al.*, 1983; Zand *et al.*, 2011). The 1° and 2° cells undergo cell divisions and morphogenic changes characteristic of their respective cell fates to produce a functioning vulva. The remaining three uninduced VPCs fuse with the surrounding hypodermis after one round of cell division (Sternberg and Horvitz, 1986).

**Figure 1.**
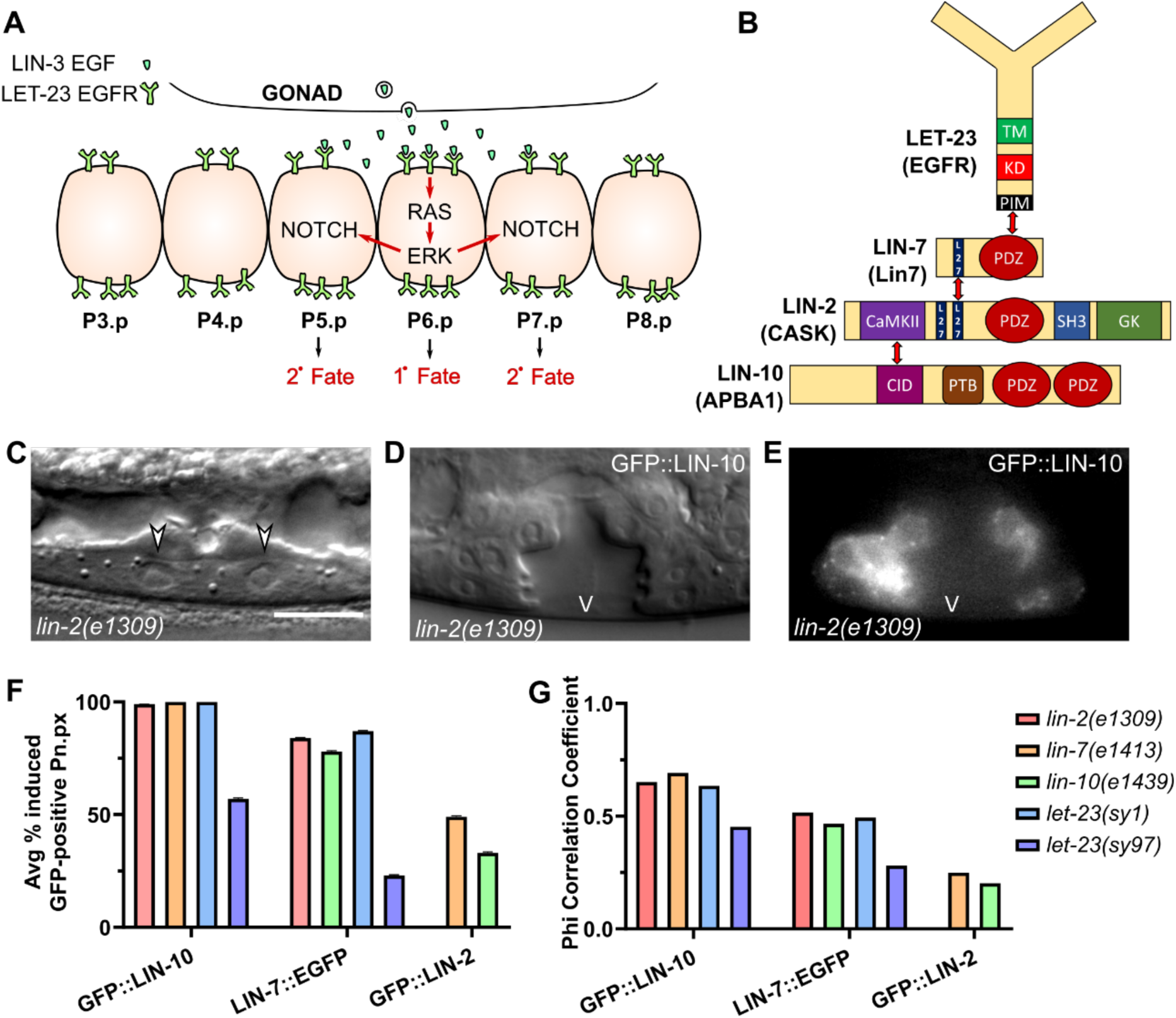
LIN-10 and LIN-7 promote VPC cell fate induction independently of their complex components. **(A)** Model of vulval cell fate induction in the vulva precursor cells (VPCs). The VPC closest to the source of the LIN-3 EGF-like ligand, P6.p, receives sufficient LET-23 EGFR activation on the basolateral membrane to activate the RAS/ERK signaling cascade, which induces the primary (1°) cell fate, and initiates lateral signaling to adjacent P5.p and P7.p through Notch to induce the secondary (2°) vulval cell fate in those cells. **(B)** Known interactions and domains between LET-23 EGFR and the LIN-2/7/10 complex in *C. elegans*. The PDZ (PSD95)/Dlg1/ZO-1) domain of LIN-7 interacts with the C-terminal PDZ-interaction motif (PIM) of LET-23 EGFR. LIN-7 and LIN-2 interact through shared L27 (LIN-2/LIN-7) domains. The CaMKII (Calcium/Calmodulin Kinase II) domain of LIN-2 interacts with the CID (CASK-interacting domain) of LIN-10. TM: Transmembrane domain. SH3: Src homology domain 3. GK: Guanylate kinase domain. PTB: Phosphotyrosine-binding domain. **(C)** Differential Interference Contrast (DIC) image of a vulvaless *lin-2(e1309)* mutant with arrows marking uninduced P6.p daughter cells. Scalebar: 10 µm. **(D-E)** DIC and epifluorescence images of an induced vulva (the lumen marked with “V”) of a *lin-2* mutant overexpressing LIN-10a (*vhEx37*). Scale as in (C). **(F)** Average percentage of cells (Pn.px lineages) expressing the indicated extrachromosomal arrays that are induced into vulval cell fate lineages. Error bars are s.d. Legend as in (G). Sample sizes for corresponding genotypes in Tables 1 and 2. **(G)** Phi correlation of extrachromosomal array expression and vulval cell fate induction for the indicated transgenes and mutant backgrounds. Sample sizes for corresponding genotypes in Tables 1 and 2.

Basolateral localization of LET-23 EGFR, which is indispensable for signaling activation, is mediated by a protein complex consisting of three PDZ (PSD95/Dlg1/ZO-1) protein interaction domain-containing scaffold proteins: LIN-2, LIN-7, and LIN-10 (Figure 1B). Named for their involvement in the vulval cell fate *lin*eage, these three proteins are essential for vulval development: loss of any individual complex component, or loss of the interaction between the C-terminal PDZ interaction motif of LET-23 EGFR to the PDZ domain of LIN-7 results in exclusive apical receptor localization and no vulval development (Ferguson and Horvitz, 1985; Simske *et al.*, 1996; Kaech *et al.*, 1998). This developmental defect prevents the worms from laying eggs, producing a striking “bag of worms” phenotype as the eggs accumulate and hatch inside the vulvaless (Vul) mother. Mammalian Lin7A-C, also known as Veli1-3 or Mals1-3, also interact with the C-terminal cytoplasmic tail of EGFR paralogs ErbB2 and ErbB4 to regulate their basolateral localization (Shelly *et al.*, 2003); however, involvement of the other complex components in receptor localization has not been determined.

The LIN-2/LIN-7/LIN-10 (LIN-2/7/10) complex has been well-defined biochemically: LIN-7 and LIN-2 interact through N-terminal L27 domains on each protein, and the calcium/calmodulin kinase II (CaMKII) domain of LIN-2 interacts with the CASK-interacting domain (CID) of LIN-10 (Figure 1B) (Borg *et al.*, 1998; Butz *et al.*, 1998; Kaech *et al.*, 1998). Still, little is known about how the complex regulates polarized LET-23 EGFR localization. LIN-2 also has a second L27 domain, which has been shown to interact with EPS-8 to prevent the internalization of LET-23 EGFR (Stetak *et al.*, 2006). Similar to other membrane-associated guanylate kinase (MAGUK) scaffold proteins, LIN-2 has a type II PDZ domain (defined by specificity for consensus interaction motifs (Songyang *et al.*, 1997)), a Src homology 3 (SH3) domain, and a C-terminal guanylate kinase domain, though these domains have not been shown to be important for vulval development. LIN-10 has a phosphotyrosine binding domain (PTB), and two tandem PDZ domains on its C-terminus with unique consensus interaction motifs that most closely resemble type III PDZ domains (Long *et al.*, 2005; Swistowski *et al.*, 2009). The PTB domain, but not the PDZ domains, was previously found to be involved in vulval development through an unknown mechanism (Whitfield *et al.*, 1999).

In addition to vulval cell fate induction, the LIN-2/7/10 complex has been found to regulate infection sensitivity in the hypodermis through an interaction between the PDZ domain of LIN-2 and the insulin receptor DAF-2 in *C. elegans* (Sem *et al.*, 2012). In mammals, the homologous CASK (LIN-2)/Lin7/APBA1 (LIN-10) complex has been well-studied for its role in regulating synaptic localization of the NMDA receptor subunit NR2B (Jo *et al.*, 1999; Setou *et al.*, 2000) and the cellular adhesion protein neurexin (Butz *et al.*, 1998; Fairless *et al.*, 2008). Lin7 is often implicated in polarized membrane sorting of internalized target proteins (Perego *et al.*, 1999; Straight *et al.*, 2001; Shelly *et al.*, 2003; Alewine *et al.*, 2007), and along with CASK, typically localizes to the basolateral membrane in epithelial cells (Cohen *et al.*, 1998; Straight *et al.*, 2000). On the other hand, Golgi-associated APBA1, also known as Mint1 or X11α, can regulate protein trafficking by interacting with the neuron-specific kinesin KIF17 (Setou *et al.*, 2000). APBA1 can also regulate secretion by interacting with the SNARE-interacting protein Munc18 (Okamoto and Sudhof, 1997; Biederer and Sudhof, 2000; Schutz *et al.*, 2005). However, this interaction is likely not conserved in *C. elegans* due to an absence of a Munc18-interacting domain in LIN-10. Furthermore, the *C. elegans* homologs of KIF17 and Munc18 (OSM-3 and UNC-18, respectively) are restricted to neurons (Gengyo-Ando *et al.*, 1993; Tabish *et al.*, 1995); therefore, they are unlikely to be involved in LIN-2/7/10 complex function in epithelial cells.

Recently, we found that LIN-10 colocalizes with LIN-2, LIN-7, and LET-23 EGFR on cytosolic punctae, likely representing Golgi ministacks and/or recycling endosomes, consistent with a model in which the complex helps target the receptor to the basolateral membrane of the VPCs. LIN-10 is strongly punctate while LIN-2 and LIN-7 are largely cytosolic, but are recruited to punctae by LIN-10. We also found that LIN-2 colocalized and interacted strongly with LIN-7 and relatively weakly with LIN-10 *in vivo*, further highlighting unique features of LIN-10 in the complex (Gauthier and Rocheleau, 2020).

Here, we report that overexpression of LIN-10 or LIN-7 can compensate for loss of their fellow complex components and restore vulval development in Vul mutants, pointing to a novel complex-independent function for both proteins. This effect is independent of the LET-23 EGFR C-terminal PDZ interaction motif, but is LET-23 EGFR-dependent. We also found a differential requirement between the PTB and PDZ domains of LIN-10 in mediating its function and localization, suggesting LIN-10 can work through a secondary pathway to promote LET-23 EGFR signaling and vulval development. These studies reveal novel contributions of LIN-10 and LIN-7 in regulating polarized protein sorting in epithelial cells.

## Results

### LIN-10 and LIN-7 can promote VPC induction in a complex-independent manner

Loss of either *lin-2, lin-7*, or *lin-10* alone has previously been shown to inhibit vulval cell fate induction due to the mislocalization of LET-23 EGFR, likely leading to a loss of signaling. Recently, we tested if loss of these genes would also disrupt localization of their complex components (Gauthier and Rocheleau, 2020). In passing, we noticed an unusually high proportion of egg-laying competent *lin-2(e1309)* mutant worms expressing a GFP-tagged LIN-10 transgene, which contrasted with *lin-2* mutants lacking the transgene that are normally Vul and egg laying-defective. This led us to suspect that there may be some compensatory mechanisms among members of the LIN-2/7/10 complex.

To test this, we scored vulval development in mutant L4 worms in which VPC lineages expressed extrachromosomal LIN-2, LIN-7, or LIN-10 transgenes that rescue their respective mutants (Gauthier and Rocheleau, 2020). These transgenes are inherently overexpressed due to expression of their respective endogenous genes, and are expressed mosaically among cells. We found that overexpression of GFP::LIN-10a (*vhEx37*) and mCherry::LIN-10a (*vhEx63*) strongly rescued VPC induction in *lin-2* and *lin-7(e1413)* null mutants without inducing defects in vulval development when overexpressed in wild type worms (Figure 1C-E; Table 1 rows 1, 4, 5, and 8). Overexpression of LIN-7a::EGFP (*vhEx60*) was also able to partially rescue vulval development in *lin-2* and *lin-10(e1439)* mutants (Table 1 rows 3 and 11), whereas overexpression of GFP::LIN-2a (*vhEx58*) failed to rescue *lin-7* or *lin-10* (Table 1 rows 7 and 10).

**Table 1.** LIN-10 and LIN-7, but not LIN-2, can promote vulval cell fate induction independently of their complex components.

|  | Genotype | %Vul | %Muv | Avg. # of VPCs induced | n |
| --- | --- | --- | --- | --- | --- |
|  | Wild type (N2) | 0% | 0% | 3.00 | many |
| 1 | *GFP::LIN-10a* | 0% | 0% | 3.00 | 17 |
| 2 | *lin-2(e1309) †* | 96% | 0% | 0.39 | 23 |
| 3 | *lin-2(e1309); LIN-7a::EGFP* | 70%* | 0% | 1.88**** | 30 |
| 4 | *lin-2(e1309); GFP::LIN-10a* | 16%**** | 8% | 2.85**** | 37 |
| 5 | *lin-2(e1309); mCh::LIN-10a ‡* | 78% | 0% | 1.30** | 27 |
| 6 | *lin-7(e1413)* | 78% | 2% | 1.15 | 51 |
| 7 | *lin-7(e1413); GFP::LIN-2a* | 79% | 0% | 1.42 | 33 |
| 8 | *lin-7(e1413); GFP::LIN-10a* | 23%**** | 13% | 2.80**** | 30 |
| 9 | *lin-10(e1439)* | 91% | 0% | 0.80 | 22 |
| 10 | *lin-10(e1439); GFP::LIN-2a* | 86% | 0% | 0.90 | 21 |
| 11 | *lin-10(e1439); LIN-7a::EGFP* | 59%* | 3% | 2.03*** | 29 |

\*p<0.05; \*\*p<0.01; \*\*\*p<0.001; \*\*\*\*p<0.0001. Fisher's exact test (%Muv and %Vul); One-way ANOVA with multiple comparisons (Avg. # of VPCs induced). Rows 3-5 were compared with row 2; Rows 7 and 8 were compared with row 6; Rows 10 and 11 were compared with row 9. †Control data collected with *lin-2(e1309); vhEx58(plin-31::gfp::lin-2a)* as published in (Gauthier and Rocheleau, 2020). ‡VPC induction scored in worms with *Pttx-3::GFP* expression because mCherry signal undetectable in VPCs under epifluorescence.

Virtually every Pn.px lineage with detectable GFP::LIN-10 transgene was induced to adopt vulval cell fates in *lin-2* and *lin-7* mutants, suggesting a strong association between LIN-10 overexpression and cell fate induction, and suggesting that LIN-10 is working cell autonomously (Figure 1F). This is supported by relatively strong Phi correlation coefficients of 0.65 and 0.69 in *lin-2* and *lin-7* mutants, respectively (Figure 1G). Correlation was more moderate for LIN-7 overexpression, with coefficients of 0.52 and 0.47 in *lin-2* and *lin-10* mutants, and with roughly 80% of LIN-7-overexpressing Pn.px lineages induced (Figure 1G-G). On the other hand, LIN-2 overexpression had minimal correlation with VPC cell fate induction (Figure 1F-G), further indicating that induction is specific to LIN-10 and LIN-7 overexpression, and not an artefact of GFP or transgene expression.

### LIN-10 and LIN-7 overexpression rescue *let-23(sy1)*, but not signaling-defective *let-23(sy97)*

LIN-7 interacts with LET-23 EGFR via its C-terminal type I PDZ domain. LIN-10 has two PDZ domains that have previously been found to recognize distinct consensus sequences (Swistowski *et al.*, 2009). Nevertheless, there is some overlap in interacting networks between different PDZ domains. The second PDZ domain of mammalian LIN-10 homolog APBA1 typically recognizes X–Φ–V/L/I motifs (where Φ is a hydrophobic residue), although it can also interact with polar residues at the −1 position (Swistowski *et al.*, 2009). Therefore, the PDZ-interaction motif of LET-23 EGFR (-TCL) is a potential match for the second PDZ domain of LIN-10.

To check if LIN-10 overexpression might independently promote signaling by interacting with LET-23 EGFR via its own PDZ domains, we tested for rescue of the *sy1* allele of *let-23* that has an early stop codon resulting in a truncation of its C-terminal PDZ interaction motif (Aroian *et al.*, 1994; Kaech *et al.*, 1998). This allele is otherwise functional for *let-23* signaling events and only causes abnormalities in vulval development (Aroian *et al.*, 1994), where interaction with LIN-7 is necessary for receptor localization and function. LIN-10 overexpression strongly rescues a *let-23(sy1)* mutant (Table 2), suggesting that LIN-10 does not require an interaction between the receptor and LIN-7 to promote vulval development, nor is it likely that the PDZ domains of LIN-10 interact with the PDZ interaction motif of LET-23 EGFR. LIN-10 expression was once again strongly correlated to cell fate induction (Figure 1F-G). Unexpectedly, LIN-7 overexpression also partially rescued the *let-23(sy1)* allele (Table 2; Figure 1F-G), suggesting that LIN-7 can also promote vulval induction independently of its interaction with the LET-23 EGFR PDZ interaction motif.

**Table 2.** LIN-10 and LIN-7 overexpression rescue a PDZ interaction-deficient *let-23(sy1)* receptor mutant, but not signaling defective *let-23(sy97)*

|  | Genotype | % Muv | % Vul | Avg. # of<br>VPCs induced | n |
| --- | --- | --- | --- | --- | --- |
| 1 | *let-23(sy1)* | 0% | 92% | 0.49 | 48 |
| 2 | *let-23(sy1); GFP::LIN-10* | 10% | 30%**** | 2.72**** | 30 |
| 3 | *let-23(sy1); LIN-7::EGFP* | 0% | 74%* | 1.94**** | 27 |
| 4 | *let-23(sy97)* | 0% | 95% | 0.48 | 43 |
| 5 | *let-23(sy97); GFP::LIN-10* | 0% | 96% | 0.72 | 25 |
| 6 | *let-23(sy97); LIN-7::EGFP* | 0% | 100% | 0.52 | 22 |

\*p<0.05; \*\*\*\*p<0.0001; Fisher's exact test (%Muv and %Vul); One-way ANOVA with multiple comparisons (Avg # VPCs induced). Rows 2 and 3 were compared with row 1; Rows 5 and 6 were compared with row 4.

To test if LIN-10 and LIN-7 overexpression might be activating signaling downstream of the receptor, we tested for rescue of a signaling-defective *let-23(sy97)* mutant that is unable to activate the downstream LET-60 Ras/MPK-1 MAPK pathway. This allele has other phenotypes associated with loss of LET-60 Ras activation, such as a rod-like lethal phenotype in L1 worms (Aroian *et al.*, 1994). Although LIN-10 overexpression moderately correlates with VPC induction in *let-23(sy97)* (Figure 1F-G), both LIN-10 and LIN-7 overexpression fail to rescue the *let-23(sy97)* allele (Table 2), suggesting that both proteins require a functioning receptor and do not promote VPC induction by activating the signaling cascade downstream of LET-23 EGFR.

### LIN-10 independently promotes basolateral LET-23 EGFR localization

The association between LIN-10 and LET-23 EGFR has previously been shown to be indirect via LIN-2 and LIN-7 (Kaech *et al.*, 1998); therefore, it is particularly surprising to find that its overexpression can promote vulval cell fate induction independently of LIN-2 and LIN-7. This suggests that LIN-10 may be working through an alternate, complex-independent pathway to promote LET-23 EGFR signaling activation.

To better understand how LIN-10 overexpression can independently promote signaling, we next tested the effect of extrachromosomal mCh::LIN-10a expression (*vhEx63*) on LET-23::GFP (*zhIs035*) localization. We found that overexpression of LIN-10 was able to restore some basolateral receptor localization in a *lin-2* mutant (Figure 2A-C). Our lab has previously reported similarly modest restoration of basolateral LET-23 EGFR localization in a *lin-2* mutant by loss of negative regulators, such as the ARF Guanine Exchange Factor AGEF-1 or the AP-1 clathrin adaptor µ-subunit UNC-101 (Skorobogata *et al.*, 2014). In a wildtype background, LIN-10 overexpression also increased the basal/apical ratio of LET-23::GFP localization (Figure 2D-F). An increase in fluorescent intensity on the basal membrane can be detected in one-cell P6.p, but not in two-cell P6.px (Figure 2G-H). This suggests that LIN-10 independently regulates LET-23 EGFR signaling by promoting basolateral receptor localization.

**Figure 2.**
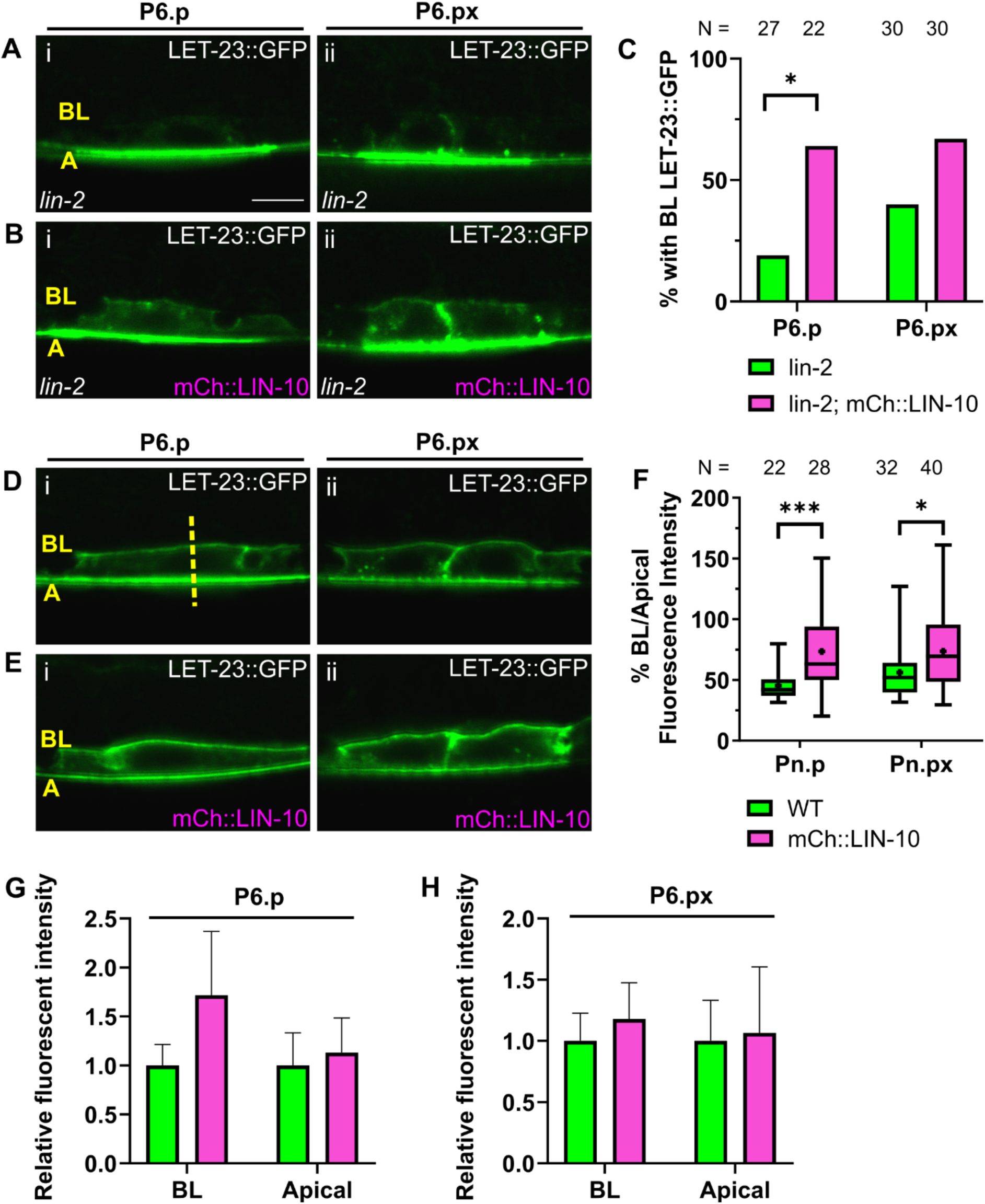
LIN-10 overexpression promotes basolateral targeting of LET-23::GFP. **(A)** LET-23::GFP (*zhIs035*) is localized on apical membranes of *lin-2(e1309)* mutant P6.p (i) and P6.px (ii) cells. **(B)** Overexpression of mCh::LIN-10 (*vhEx63*) restores basolateral LET-23::GFP in a *lin-2* mutant. **(C)** Quantification of VPCs with basolateral LET-23::GFP localization in (A) and (B). *p<0.05 Fisher’s exact test. **(D)** Polarized LET-23::GFP localization in wildtype P6.p (i) and P6.px (ii) cells. **(E)** LET-23::GFP localization in VPCs expressing extrachromosomal mCh::LIN-10. **(F)** Quantification of basolateral/apical peak fluorescent intensity of images represented in (D-E). *p<0.05, ***p<0.001 Two-tailed Student’s *t*-test. **(G-H)** Comparing relative fluorescent intensity of basolateral and apical membranes in P6.p (G) and P6.px (H) with or without LIN-10 overexpression for the set of images represented in (D-E). Legend and sample sizes as in (F). Intensities normalized to fluorescent intensity of wildtype worms (no mCh::LIN-10 expressed) for basolateral and apical membranes. Error bars are s.d. BL = Basolateral. A = Apical. WT = Wildtype. Scalebar: 5 µm. Images are all at same scale.

### LIN-10 C-terminal domains mediate punctate localization

LIN-10 localizes to Golgi ministacks and recycling endosomes in neurons and intestinal epithelia (Rongo *et al.*, 1998; Glodowski *et al.*, 2005; Zhang *et al.*, 2012), and shows strong punctate localization in the VPCs (Gauthier and Rocheleau, 2020). To better understand the function of LIN-10, we performed a structure-function analysis to identify the protein domains that regulate its localization and function (Figure 3A-B). The N-terminal half of LIN-10 has no characterized domains other than the LIN-2-interacting CID, shared only with mammalian APBA1. Overall, the N-terminus of LIN-10 shares low sequence homology with APBA1 and APBA2, who in turn share low sequence homology in their N-terminal domains with each other, despite both sharing MIDs (Figure 3A and (Glodowski *et al.*, 2005)). Despite divergent sequences, the function of the CID is conserved between LIN-10 and APBA1, as *C. elegans* LIN-10 can interact with mammalian CASK (Borg *et al.*, 1998). On the other hand, the C-terminal half, containing the PTB and tandem PDZ domains, has a much higher sequence identity among LIN-10 and APBA1-3 paralogs (Figure 3A and (Glodowski *et al.*, 2005)).

**Figure 3.**
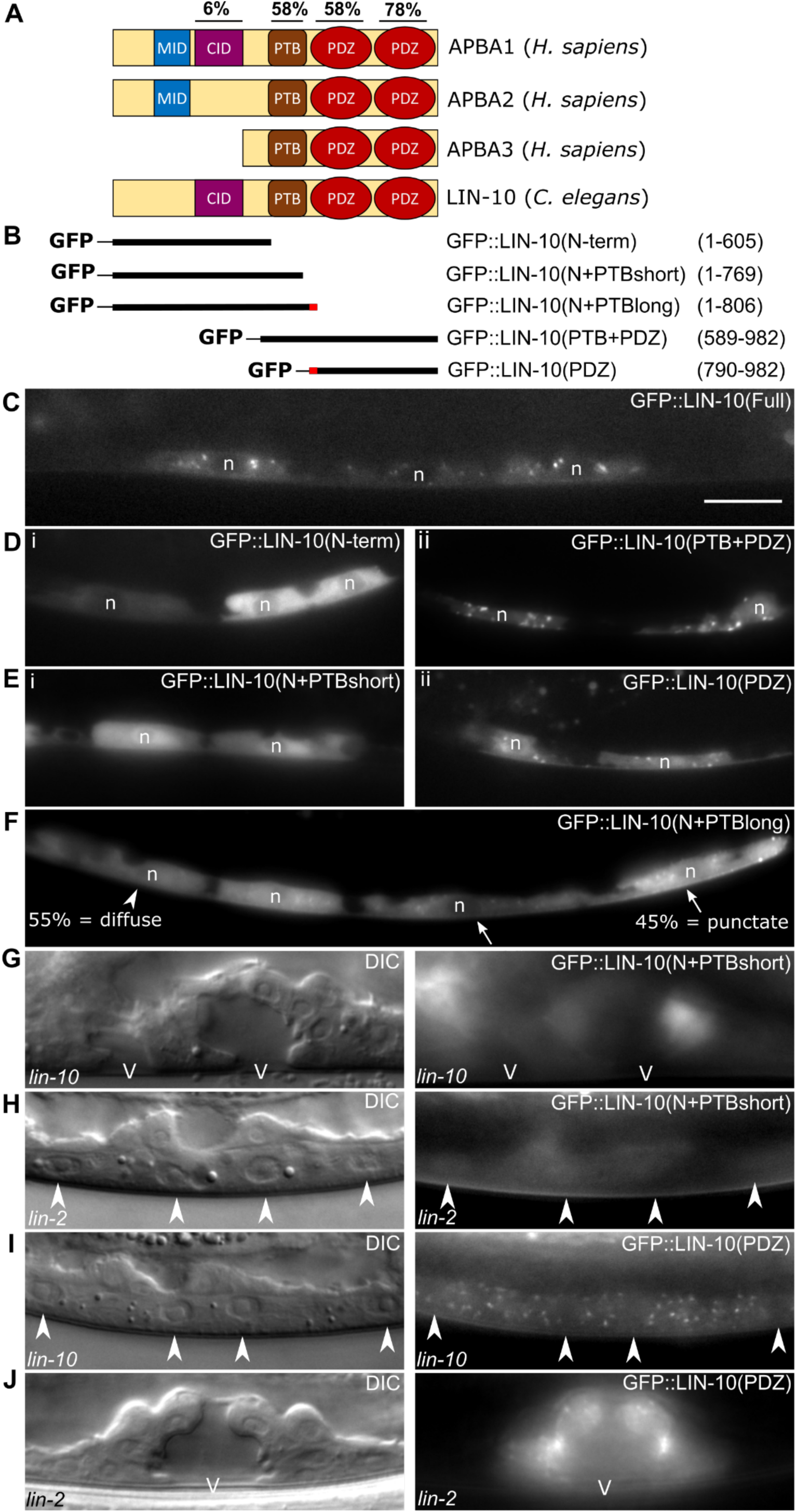
C-terminal domains regulate punctate localization and function of LIN-10. **(A)** Protein domains of human APBA1-3 and *C. elegans* LIN-10. Percent identities compare amino acid sequences of CASK Interaction Domain (CID), Phosphotyrosine Binding Domain (PTB), and PDZ1/2 domains of LIN-10a to those of APBA1. **(B)** Truncations of LIN-10a generated to compare localization and function. Red region of N+PTBlong and PDZ indicates 16 amino acids overlapping between these two truncations. **(C-F)** Localization of full-length GFP::LIN-10 (*vhEx37*) (C), and the N-term (*vhEx52*) (D i), PTB+PDZ (*vhEx55*) (D ii), N+PTBshort (*vhEx74*) (E i), PDZ (*vhEx66*) (E ii), and N+PTBlong (*vhEx72*) (F) truncated LIN-10 fragments as shown in (B). N in (F) = 29 cells imaged (from 19 worms). Arrowhead: VPC with diffuse cytosolic localization. Arrows: VPCs with punctate localization. **(G-H)** Expression of the N+PTBshort construct rescues *lin-10(e1439)* (G), but not *lin-2(e1309)* (H). A pseudovulva, and a partially induced vulva can be seen in (G). **(I-J)** Expression of the PDZ domains fails to rescue a *lin-10(e1439)* mutant (I) but does rescue a *lin-2(e1309)* mutant (J). Arrowheads in (H) and (I) are nuclei of uninduced primary (two in the centre) and secondary (on left and right edge of image) Pn.px cells. n: Nucleus. V: Vulval lumen. Scalebar: 10 µm. Images are all at same scale.

We cloned the CID-containing N-terminal half (LIN-10(N-term), aa 1-605) and the PTB and PDZ-containing C-terminal half (LIN-10(PTB+PDZ), aa 589-982) of LIN-10 to identify which region of LIN-10 regulates its localization and function (Figure 3B). Whereas full-length LIN-10 localizes to cytosolic punctae in all VPCs (Figure 3C), we found that the LIN-10(N-term) fragment was diffusely expressed in the cytosol (Figure 3D i). In contrast, the C-terminal LIN-10(PTB+PDZ) showed robust localization to punctae (Figure 3D ii), indicating that these domains mediate localization to Golgi and endosomes. Inclusion of the PTB domain in the N-terminal fragment, LIN-10(N+PTBshort) (aa 1-769) was not sufficient to localize to puncta (Figure 3E i), whereas omitting the PTB domain from the C-terminal fragment, as in LIN-10(PDZ) (aa 790-982), did not prevent punctate localization (Figure 3E ii), suggesting that the PDZ domains mediate this localization pattern. Interestingly, extending the N-terminal half beyond the PTB domain into the flexible linker region between the PTB and the first PDZ domain, as in LIN-10(N+PTBlong) (aa 1-806), recovered some punctate localization, even in a *lin-10* mutant (Figure 3F). This LIN-10(N+PTBlong) fragment overlaps with the LIN-10(PDZ) domain fragment by 16 amino acids covering a small segment of the linker and PDZ1 domain (Figure 3B), suggesting there may be a regulatory region in this segment that in part determines punctate localization with the PDZ domains.

Mammalian APBA1 have been shown to have intramolecular interactions in their PDZ1 domain (Long *et al.*, 2005), and *C. elegans* LIN-10 has been found to self-interact in a yeast two-hybrid screen (Walhout *et al.*, 2000). To verify that the localization pattern of the PDZ domains was not due to oligomerization with endogenous full-length LIN-10, we looked at its localization in a *lin-10* mutant, and still found that the PDZ domains were punctate (Figure 3I). Therefore, these results demonstrate that the C-terminal PDZ domains, and likely a portion of the linker region, mediate Golgi and endosomal localization of LIN-10.

### C-terminal PDZ domains mediate complex-independent function of LIN-10, whereas PTB domain required for overall LIN-10 function

To further characterize the complex-independent function of LIN-10, we tested which domains of LIN-10 were necessary for the rescue of *lin-2* and *lin-10*. We found that the C-terminal PDZ domains that mediated punctate localization of LIN-10 were necessary and sufficient to rescue VPC induction in a *lin-2* mutant (Table 3 rows 8, 9, and 11; Figure 3H, J). However, expression of these domains alone was not sufficient to rescue *lin-10* (Table 3 row 6; Figure 3I). On the other hand, while truncation of the PDZ domains in the LIN-10(N+PTBshort) fragment failed to rescue a *lin-2* mutant, this fragment did rescue *lin-10* (Table 3 rows 3 and 9; Figure 3G-H). We found that presence of the PTB domain, whether on the N-terminal cytosolic half (N+PTBshort or N+PTBlong) or added to the C-terminal punctate domains (PTB+PDZ), was specifically required for rescue of a *lin-10* mutant. However, *lin-10* rescue by the LIN-10(PTB+PDZ) fragment was mild compared to the N-terminal fragments, consistent with the CID-mediated interaction with LIN-2 being important for LIN-10 complex-dependent function (Table 3 rows 3-5). Furthermore, LIN-10(N+PTBlong) rescue is stronger than N+PTBshort, suggesting that Golgi and endosome localization is functionally important (Table 3 rows 3-4). Interestingly, LIN-10(PDZ) rescues *lin-2* Vul phenotype slightly better than LIN-10(PTB+PDZ), suggesting that the PTB domain might interfere with the PDZ-mediated *lin-2* rescue (Table 3 rows 10-11). These results suggest that the PTB and PDZ domains are differentially required for LIN-10 function: general LIN-10 function requires the PTB domain, and is less dependent on the subcellular localization, whereas the LIN-2-independent function of LIN-10 is likely restricted to Golgi and endosomal subcellular compartments and is mediated by the C-terminal PDZ domains.

**Table 3.** Differential requirements of the LIN-10 PTB and PDZ domains for rescue of *lin-2* and *lin-10* mutants.

|  | Genotype | %Vul | %Muv | Avg # of VPCs induced | n |
| --- | --- | --- | --- | --- | --- |
| 1 | *lin-10(e1439)* | 90% | 0% | 0.86 | 49 |
| 2 | *lin-10(e1439); GFP::LIN-10(N-term)* | 91% | 3% | 1.27 | 35 |
| 3 | *lin-10(e1439); GFP::LIN-10(N+PTBshort)* | 52%**** | 11%* | 2.39**** | 27 |
| 4 | *lin-10(e1439); GFP::LIN-10(N+PTBlong)* | 4%*** | 4% | 3.00**** † | 24 |
| 5 | *lin-10(e1439); GFP::LIN-10(PTB+PDZ)* | 77% | 3% | 1.85**** | 30 |
| 6 | *lin-10(e1439); GFP::LIN-10(PDZ)* | 95% | 0% | 0.73 | 22 |
| 7 | *lin-2(e1309)* | 88% | 0% | 0.64 | 60 |
| 8 | *lin-2(e1309); GFP::LIN-10(N-term)* | 81% | 0% | 1.08 | 31 |
| 9 | *lin-2(e1309); GFP::LIN-10(N+PTBshort)* | 100% | 0% | 0.62 | 26 |
| 10 | *lin-2(e1309); GFP::LIN-10(PTB+PDZ)* | 74% | 0% | 1.73**** | 31 |
| 11 | *lin-2(e1309); GFP::LIN-10(PDZ)* | 54%*** | 4% | 2.25**** | 24 |

\*p<0.05; \*\*\*p<0.001; \*\*\*\*p<0.0001; Fisher's exact test (%Muv and %Vul); One-way ANOVA with multiple comparisons (Avg # VPC induced). Rows 2 to 6 were compared with row 1. Rows 8 to 11 were compared to row 7. †p<0.01 Student's two-tailed *t*-test comparing row 4 to row 3.

### ARF-1.2 colocalizes with LIN-10, but is not required for LIN-10 overexpression rescue of *lin-2*

The finding that the C-terminal PDZ domains of LIN-10 mediate its complex-independent function suggests that LIN-10 has additional binding partners with which it promotes basolateral LET-23 EGFR targeting and signaling. The C-terminus (PDZ domains and part of the PTB domain) of all three mammalian APBA proteins mediate an interaction with GTP-bound Class I (Arf 3) and Class II (Arf4) Arf GTPases in neurons to regulate Amyloid Precursor Protein (APP) trafficking (Hill *et al.*, 2003). APBA proteins were also found to localize to the Golgi in a Brefeldin A (BFA)-sensitive manner, a compound that disrupts the Arf guanine exchange factors (GEF) BIG1/2 and GBF1 (Donaldson *et al.*, 1992a; Donaldson *et al.*, 1992b; Togawa *et al.*, 1999; Niu *et al.*, 2005), and are associated with clathrin-coated secretory vesicles (Hill *et al.*, 2003; Shrivastava-Ranjan *et al.*, 2008). Our lab has previously reported that the *C. elegans* Class I and Class II Arf GTPases (ARF-1.2 and ARF-3, respectively), work in opposition to the LIN-2/7/10 complex with the Arf GEF AGEF-1 (homologous to BIG1/2) and the AP-1 clathrin adaptor complex to negatively regulate LET-23 EGFR signaling and basolateral localization (Skorobogata *et al.*, 2014), possibly by promoting the apical targeting of the receptor. LIN-10 overexpression phenocopies inhibition of the AGEF-1/ARF/AP-1 pathway with respect to rescue of VPC induction and increasing basolateral LET-23 EGFR localization. Therefore, the complex-independent function of LIN-10 might involve an interaction with ARF GTPases that could inhibit or compete with the AP-1-mediated pathway.

ARF-1.2::EGFP localizes to cytoplasmic foci when expressed in the VPCs (O Skorobogata and CER, unpublished results). We found that LIN-10 frequently colocalizes with ARF-1.2 cytoplasmic foci or can localize to adjacent foci at Pn.p and Pn.px stages (Figure 4A-B). Unlike in mammals, LIN-10 does not require active ARF GTPases for its localization, as punctate localization is not disrupted in *arf-1.2* loss-of-function mutants, *arf-1.2; arf-3(RNAi)*-treated mutants, or *agef-1(vh4)* hypomorphic mutants (Figure 4C). Similarly, loss of *lin-10* does not disrupt recruitment of ARF-1.2 to cytosolic punctae (Figure 4D). If ARF-1.2 had a dual role in regulating LET-23 EGFR trafficking by either antagonizing basolateral targeting when associated with AP-1, or promoting basolateral targeting when associated with LIN-10, we would expect loss of *arf-1.2* to suppress the rescue of *lin-2* by LIN-10 overexpression. Loss of *arf-1.2* only partly rescues *lin-2* (Skorobogata *et al.*, 2014), and the level of rescue is significantly less than when LIN-10 is overexpressed (Table 4). However, we found no change in vulval induction in *arf-1.2; lin-2; GFP::LIN-10* compared to *lin-2; GFP::LIN-10* (Table 4). Therefore, LIN-10 is not likely working as an ARF-1.2 effector to promote basolateral LET-23 EGFR trafficking. However, it remains possible that LIN-10 overexpression may instead function through inhibition of ARFs.

**Table 4.** ARF-1.2 not required for the rescue of *lin-2* by LIN-10 overexpression.

|  | Genotype | %Vul | %Muv | Avg # of VPCs induced | n |
| --- | --- | --- | --- | --- | --- |
| 1 | *lin-2(e1309); GFP::LIN-10a †* | 16% | 8% | 2.85 | 37 |
| 2 | *arf-1.2(ok796); lin-2(e1309)* | 85% | 5% | 1.88**** | 20 |
| 3 | *arf-1.2(ok796); lin-2(e1309); GFP::LIN-10a* | 24% | 12% | 2.94 | 17 |

\*\*\*\*p<0.0001. Fisher's exact test (%Muv and %Vul); One-way ANOVA with multiple comparisons (Avg # VPCs induced). Rows 2 and 3 were compared with row 1. †Data collected along with that in Table 1.

**Figure 4.**
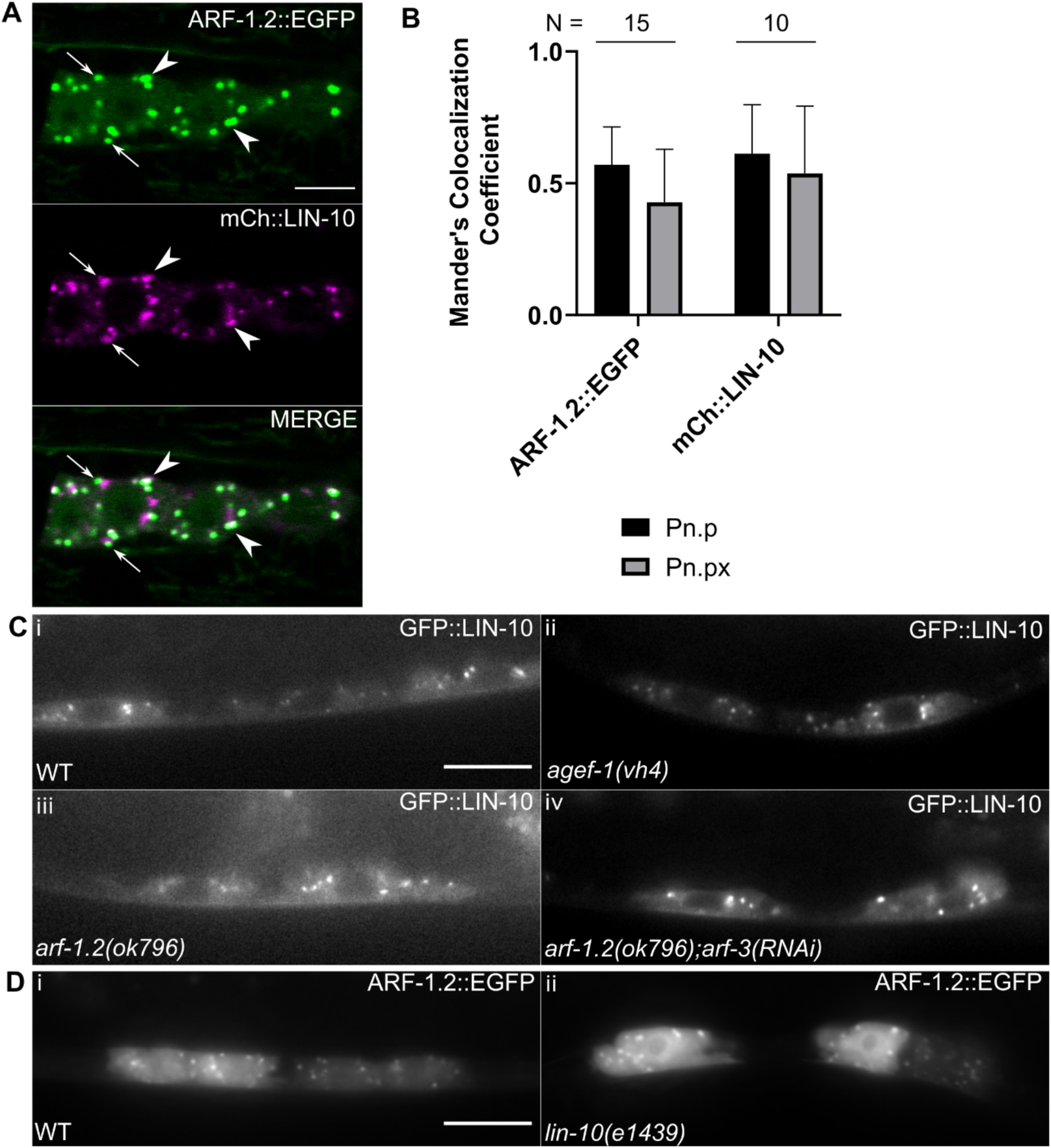
LIN-10 colocalizes with ARF-1.2 but is not dependent on ARFs for localization. **(A)** LIN-10 (*vhEx63*) frequently localizes with (arrowheads) or near (arrows) ARF-1.2 GTPase (*vhEx5*). Scalebar: 5 µm. **(B)** Average Mander’s correlation coefficients reveal moderate colocalization between ARF-1.2 and LIN-10 in *C. elegans* VPCs. Error bars are s.d. **(C)** LIN-10 (*vhEx37*) localization to cytosolic punctae (i) is not altered in a hypomorphic *agef-1(vh4)* mutant (ii), a loss-of-function *arf-1.2(ok796)* mutant (iii), and an *arf-1.2* mutant treated with *arf-3(RNAi)*. **(D)** ARF-1.2 (*vhEx7*) localization to cytosolic punctae (i) is not altered in a loss-of-function *lin-10(e1439)* mutant. Scalebars: 10 µm.

## Discussion

Here we identified a novel, complex-independent function for LIN-7 and LIN-10 in promoting LET-23 EGFR signaling and vulval cell fate induction. Overexpression of LIN-7 and LIN-10 rescued vulval development in *lin-2/10* and *lin-2/7* mutants, respectively, as well as in *let-23(sy1)* mutants deficient for an interaction with the LIN-7 PDZ domain. Focusing on LIN-10, the most removed from LET-23 EGFR, we found that it promoted basolateral receptor targeting even in absence of its complex components. We found that the LIN-10 PTB domain, but not the PDZ domains, was required for the rescue of *lin-10* mutants. Conversely, we found that the complex-independent function of LIN-10 required the C-terminal PDZ domains, that are largely responsible for its punctate localization, and not the PTB domain. Finally, LIN-10 colocalizes with ARF-1.2 in the VPCs, but does not require ARF-1.2 for localization or for its complex-independent function.

Previous research on LIN-2, LIN-7, and LIN-10 have led towards a model in which these proteins function *in vivo* as a complex, or at least as interacting components of a linear pathway (Kaech *et al.*, 1998). Therefore, we were surprised to find that overexpression of either LIN-7 or LIN-10 can compensate for loss of the other components. This suggests that our assumptions about how the complex functions are incorrect, or that LIN-7 and LIN-10 can promote LET-23 EGFR signaling via complex-independent mechanisms. Overexpression of LIN-7 can rescue the Vul phenotypes of *lin-2* and *lin-10* mutants, but not as strong as rescue of *lin-7* mutants (Gauthier and Rocheleau, 2020), suggesting that compensation for loss of LIN-2 and LIN-10 is partial. Since LIN-7 interacts directly with LET-23 EGFR, it is perhaps not too surprising that increased levels of LIN-7 would rescue *lin-2* or *lin-10* mutants. Mammalian Lin7 is involved in recycling of internalized cargo, including the EGFR paralog ErbB2, at the basolateral membrane, with no known involvement of mammalian LIN-2 or LIN-10 (CASK and APBA1-3, respectively) (Perego *et al.*, 1999; Shelly *et al.*, 2003); therefore, LIN-7 might similarly promote recycling of a small amount of LET-23 EGFR that makes it to the basolateral membrane in the absence of LIN-2 or LIN-10.

We recently found that LIN-7, but not LIN-2 or LIN-10, colocalizes with LET-23 EGFR on basolateral membranes of VPCs (Gauthier and Rocheleau, 2020), consistent with LIN-7 potentially having uncharacterized complex-independent roles in basolateral membrane traffic. Furthermore, LIN-7 basolateral localization was not dependent on the LET-23 EGFR PDZ interaction motif (Gauthier and Rocheleau, 2020). Similarly, we found that LIN-7 overexpression rescues a *let-23(sy1)* receptor mutant lacking a PDZ interaction motif, suggesting that LIN-7 can promote signaling independently of this interaction. Mammalian Lin7A was found to have a second point of interaction with the kinase domain of all four human EGFR paralogs through its extended N-terminus that helps transit the receptor from the ER to the Golgi (Shelly *et al.*, 2003). Although *C. elegans* LIN-7 only has a portion of this N-terminal sequence (Butz *et al.*, 1998; Shelly *et al.*, 2003), a similar interaction could explain how LIN-7 overexpression may rescue *let-23(sy1)* by upregulating LET-23 EGFR secretion (Figure 5). Alternatively, LIN-7 orthologs are also known to maintain epithelial cell polarity through L27 and PDZ domain-mediated interactions with the Crumbs complex and the cadherin-catenin complex at cell junctions (Perego *et al.*, 2000; Straight *et al.*, 2006). Thus, LIN-7 overexpression in the VPCs might have more global impacts on epithelial cell polarity that could indirectly shift the balance between apical and basolateral cargo sorting (Figure 5), thereby restoring some LET-23 EGFR signaling activation.

**Figure 5.**
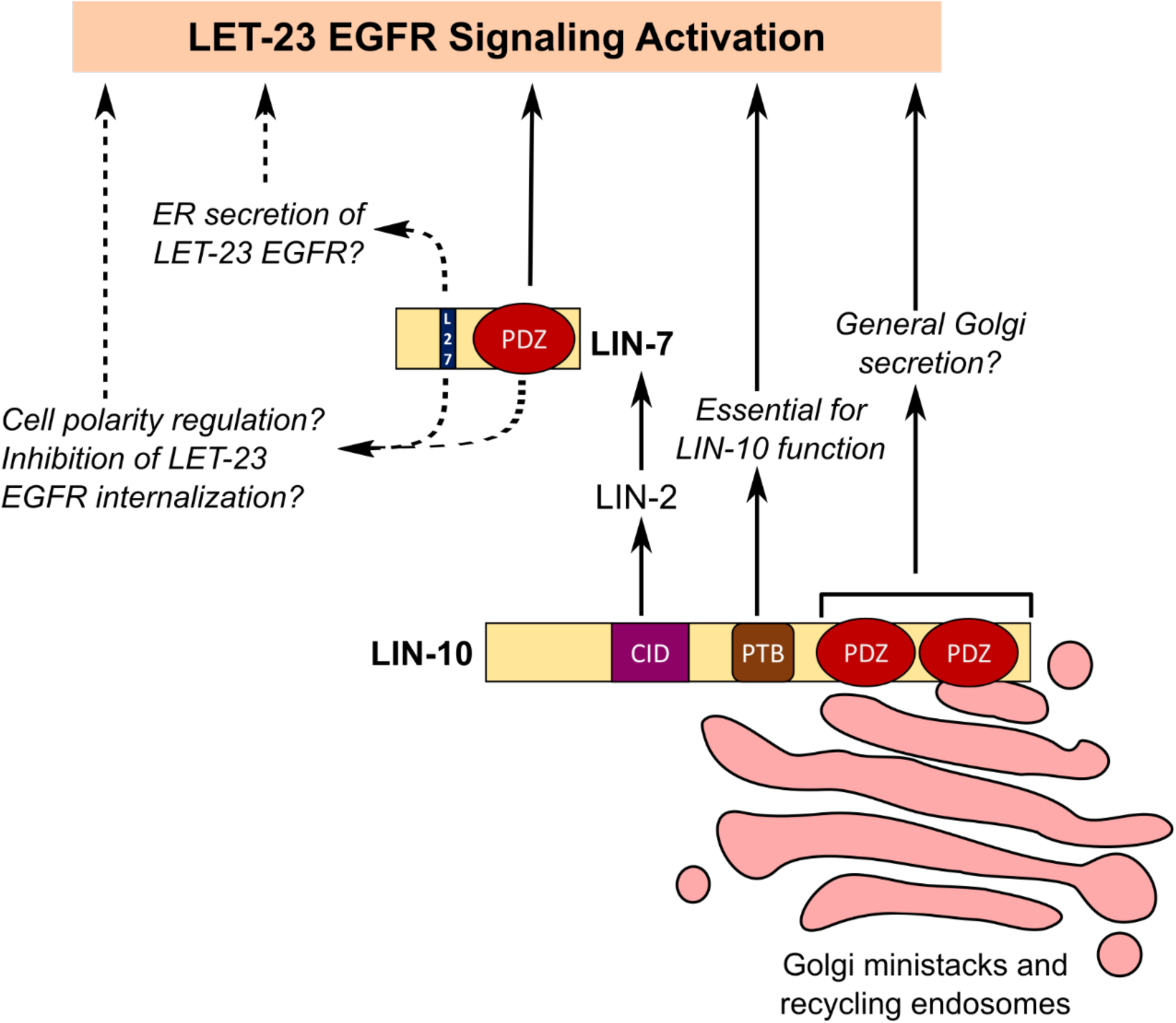
Hypothetical models for complex-independent function of LIN-7 and LIN-10 in promoting LET-23 EGFR signaling. LIN-10 targets LET-23 EGFR to the basolateral membrane through LIN-2 and LIN-7, and requires its PTB domain for its function in the VPCs. However, overexpression of its PDZ domains can bypass the requirement for its complex to promote LET-23 EGFR signaling, possibly through general upregulation of Golgi secretion. LIN-7 was also found to have a complex-independent function (dashed arrows). This may involve promoting the secretion of LET-23 EGFR from the ER (endoplasmic reticulum), inhibiting LET-23 EGFR internalization, or regulating cell polarity, similar to its mammalian orthologs.

LIN-2 is the only member of the complex that has been shown to have an additional binding partner in EPS-8 to regulate LET-23 EGFR trafficking (Stetak *et al.*, 2006). While overexpression of LIN-2 strongly rescues a *lin-*2 mutant (Gauthier and Rocheleau, 2020), it could not bypass its complex components to activate signaling. The strong binding, colocalization, and similar expression dynamics between LIN-2 and LIN-7 (Gauthier and Rocheleau, 2020) may point to a requirement for some dependence on stoichiometry with LIN-7 for LIN-2 function.

The finding that LIN-10 overexpression can rescue *lin-2* and *lin-7* mutants and partly restore LET-23 EGFR localization to the basolateral membrane of the VPCs was particularly unexpected since its interaction with LET-23 EGFR is mediated by LIN-2 and LIN-7 (Kaech *et al.*, 1998). One intriguing possibility was that the PDZ domains of LIN-10 might interact directly with the PDZ interaction motif of LET-23 EGFR. Consistent with previous protein interaction assays that did not find a direct interaction between LIN-10 and LET-23 EGFR (Kaech *et al.*, 1998), we found that LIN-10 overexpression was able to rescue the *let-23(sy1)* mutation lacking the PDZ interaction motif, suggesting that LIN-10 was not promoting LET-23 localization via a direction interaction with its PDZ domains.

The mammalian homologs of LIN-10 (APBA1-3) have well-established roles in regulating secretion and recycling. APBA2/3 lack the CASK/LIN-2-interacting CID, indicating that APBA proteins do not rely on CASK and mLin7 for their function. In *C. elegans* neurons, LIN-10 regulates the trafficking and recycling of the glutamate receptor GLR-1 independent of LIN-2 and LIN-7, and without a direct interaction to GLR-1 (Rongo *et al.*, 1998). In general, LIN-10 has relatively weak colocalization and *in vivo* association with LIN-2 and LIN-7 (Gauthier and Rocheleau, 2020). In the VPCs, LIN-2 and LIN-7 show dynamic changes in expression and localization not shared by LIN-10 (Gauthier and Rocheleau, 2020), consistent with LIN-10 potentially having complex-independent functions.

We found differential requirements for the PTB and PDZ domains for rescue of *lin-10* and *lin-2* mutants, respectively. A previous study by Glodowski, Wright, Martinowich et al., queried which domains of LIN-10 were required for GLR-1 localization in neurons and vulva induction by testing for rescue of associated *lin-10* phenotypes. They found that the PTB domain is essential for *lin-10* rescue in both tissues, whereas the PDZ domains were essential for rescue of GLR-1 localization in neurons, but not for vulval development (Glodowski *et al.*, 2005). Our results support their findings that the PTB domain is required for rescue of the *lin-10* Vul phenotype (Figure 5), and that the PDZ domains are not. We found that the PTB domain, whether part of the LIN-10 N-terminal constructs containing the CID or part of the LIN-10 C-terminal fragment containing the PDZ domains, could increase VPC induction in *lin-10* mutants. However, only the N-terminal fragments significantly suppressed the *lin-10* mutant Vul phenotype, consistent with the N-terminal CID-mediated interaction with LIN-2 being important for LIN-10 function. How the PTB domain promotes LIN-10 function in VPC induction remains unclear. Interestingly, expression of the LIN-10 C-terminal PDZ domains alone were sufficient for rescue of *lin-2* mutants, despite being dispensable for rescue of *lin-10*.

We recently demonstrated that LIN-10 recruits LIN-2 and LIN-7 to Golgi and recycling endosomes, where it also colocalizes with LET-23 EGFR (Gauthier and Rocheleau, 2020). The LIN-10 C-terminal PDZ fragment that shows the strongest localization to these subcellular compartments, despite only being involved in the complex-independent function of LIN-10. The N+PTBshort fragment, despite not localizing to punctae, is able to rescue a *lin-10* mutant, suggesting LIN-10 function may not be entirely dependent on its endomembrane localization. Still, extending this N-terminal fragment into the linker region confers an even stronger rescue while also partially restoring some punctate localization, indicating that localization to Golgi and/or recycling endosomes is likely the primary site of LIN-10 function both with and without LIN-2 and LIN-7.

The strong punctate localization and rescue of *lin-2* by the C-terminal PDZ domain suggests that LIN-10 engages with additional interactors in the VPCs through its PDZ domains to regulate LET-23 EGFR trafficking independently of LIN-2 and LIN-7. These interactions may support general secretion from the Golgi (Figure 5), as has been hypothesized for the PDZ domains of APBA2 (Saito *et al.*, 2011). The class I and II ARF-1.2 and ARF-3 GTPases are strong candidate effectors for the PDZ-mediated complex-independent function of LIN-10. Mammalian class I and II Arf GTPases interact with and might recruit APBA proteins to the Golgi to regulate secretion of APP, and APBA proteins are hypothesized to serve as clathrin adaptor proteins (Hill *et al.*, 2003). If the interaction between APBA proteins and Arf GTPases is conserved in *C. elegans*, LIN-10 might compete with AP-1 for binding to ARF-1.2 and ARF-3 to promote basolateral targeting of LET-23 EGFR. Alternatively, an interaction between LIN-10 and the ARFs might suppress the normally inhibitory function of ARF-1.2 without directly promoting basolateral targeting. The relatively strong colocalization between LIN-10 and ARF-1.2 support such a model, despite the AGEF-1 and the ARFs not being required for LIN-10 localization. The effects of LIN-10 overexpression and *arf-1.2* mutations on vulval development were not additive, consistent with these two proteins working in the same pathway; however more studies will be required to validate this hypothesis.

Thus, these findings reveal a complex-independent role for LIN-10 in promoting LET-23 EGFR trafficking to the basolateral membrane. This role may play only a minor given that under normal expression levels, LIN-10, cannot compensate for loss of *lin-2* or *lin-7.* However, it reveals a potential mechanism, by which the PDZ domains can influence LET-23 EGFR trafficking and signaling that may be critical under different circumstances and with different cargoes.

## Materials and Methods

### Strains and maintenance

All strains were maintained at 20°C on Nematode Growth Media seeded with HB101 *Escherichia coli* (Brenner, 1974). A complete strain list can be found in Supplementary Table 1.

### Vulval cell fate induction scoring

VPC induction scores were analyzed at the L4 larval stage by counting the number of induced vulval cells derived from each VPC. Wild type worms have a VPC induction score of 3 because only 3 of 6 VPCs (P5.p-P7.p) are induced. Worms were classified as Vul if they had a VPC induction score of less than 3, or multivulva (Muv) if they had an induction score of greater than 3 (Gauthier and Rocheleau, 2017). Worms with detectable fluorescent signal in their vulval cells (driven by the *lin-31* promoter) and in neurons in their head (driven by the *ttx-3* promoter of the co-expression marker) were compared to siblings taken from the same plate with no fluorescent signal in both their vulval cells and neurons as negative controls.

### LIN-10 subdomain cloning

Subdomains for the LIN-10a isoform were mapped according to the domains previously identified for the LIN-10b isoform (Glodowski *et al.*, 2005). Primers for cloning are listed in Supplementary Table 2. Subdomains were cloned into a *plin-31::gfp::lin-10a* expression vector by replacing the full-length *lin-10a* open reading frame (Gauthier and Rocheleau, 2020). All constructs were verified by sequencing. Amino acid sequence identity between LIN-10a domains and mammalian APBA1 was calculated according to sequence identities identified by BLAST (NCBI, NIH) and Clustal Omega (EMBL-EBI). Total number of similar amino acids were divided by the total number of amino acids in the respective LIN-10 protein domain to calculate sequence identity.

### Microscopy and image analysis

Worms were immersed in 10 mM levamisole on a 2% agarose pad on a glass slide prior to imaging and scoring on an Axio A1 Imager epifluorescent microscope (Zeiss). Images were analyzed using FIJI (ImageJ). Confocal (colorized) images were taken using an LSM780 laser-scanning confocal microscope. Mander’s correlation coefficients for ARF-1.2::EGFP and mCherry::LIN-10 were measured using Zen 2012 software (Zeiss) and quantified by tracing the cells of interest, yielding a software-generated scatterplot of green and red fluorescent intensity values per pixel. Crosshairs along the X and Y axis divide the graph into four quadrants and are used to omit background fluorescence, such that only pixels in the upper right-hand quadrant are calculated for colocalization. The crosshairs were adjusted to omit most of the cytosolic signal in order to capture colocalization at punctae.

### Correlation analysis for extrachromosomal array expression and VPC induction

In addition to scoring VPC cell fate induction, we kept track of which specific Pn.px lineages expressed the extrachromosomal arrays (determined by visible GFP fluorescence on a Zeiss Axio A1 imager at 100X) in developing vulvas of early to mid-L4 larvae. For each worm, we counted the number of GFP-expressing Pn.px lineages that were induced into vulval cell fates (either the primary or secondary cell fates), or that were uninduced (did not continue dividing).

To assess the correlation between extrachromosomal array expression and VPC cell fate induction, Pn.px lineages of all worms per genotype were categorized into four groups: those that expressed GFP and that were induced to assume a vulval cell fate (A), those with no visible GFP that were induced (B), those with GFP expression that were not induced (C), and those with no visible GFP that were not induced (D). Using these groups, the Phi (Φ) Correlation Coefficient between GFP expression and VPC induction was calculated for each worm using the formula below, and the average correlation coefficient for each genotype was used for analysis. 

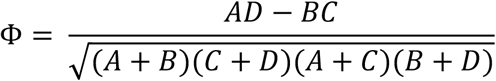

### Analyzing LET-23 EGFR polarized distribution

Polarized membrane distribution of LET-23 EGFR was analyzed by drawing a 20 pixel-wide line across P6.p cells (as in Figure 2D i) for each worm and measuring the averaged peak fluorescent intensity on the basal and apical membrane using FIJI. For worms at the 2-cell stage, one line was drawn across each P6.px cell. To limit bias, lines were drawn across DIC images over the centre of the nucleus, and transposed to the fluorescent image.

To assess restoration of basolateral LET-23 EGFR localization in *lin-2* mutants, we similarly drew a 20 pixel-wide line across the basal membrane of P6.p and P6.px cells. Cells with a peak in fluorescent intensity across the basal membrane that was at least twice as bright as the cytosol were considered to have basolateral localization of LET-23 EGFR.

### RNA interference (RNAi)

RNAi was performed by feeding as previously described (Kamath *et al.*, 2001). All RNAi experiments were performed at 20°C. RNAi clone for *arf-3* (IV-4E13) was obtained from Dr. Julie Ahringer’s RNAi library.

### Statistical analysis

Average VPC induction scores and average peak basal/apical fluorescent intensities were compared using Student’s t-test or One-way ANOVA with Dunnett’s correction for multiple comparisons. Proportions of Vul and Muv phenotypes, and proportion of P6.p and P6.px cells with basolateral LET-23 EGFR localization were compared using Fisher’s exact test. Statistical analyses were performed using GraphPad Prism 8.0.

## Acknowledgements

We thank Jung Hwa Seo, Min Fu, and ShiBo Feng for technical assistance; Dr. Richard Roy for sharing his RNAi library; and WormBase for strain information. Confocal microscopy was conducted in the Molecular Imaging Platform of the RI-MUHC. Some strains were provided by the CGC which is funded by an NIH Office of Research Infrastructure Program grant (P40 OD010440). This work was funded by an NSERC Discovery Grant (RGPIN-2018-05673) to CER. KDG was funded by Canada Graduate Scholarship Master’s Award and Post-Graduate Scholarship Doctoral fellowships from NSERC, and Master’s and Doctoral Awards from FRSQ.

---

**Supplementary Table 1:**
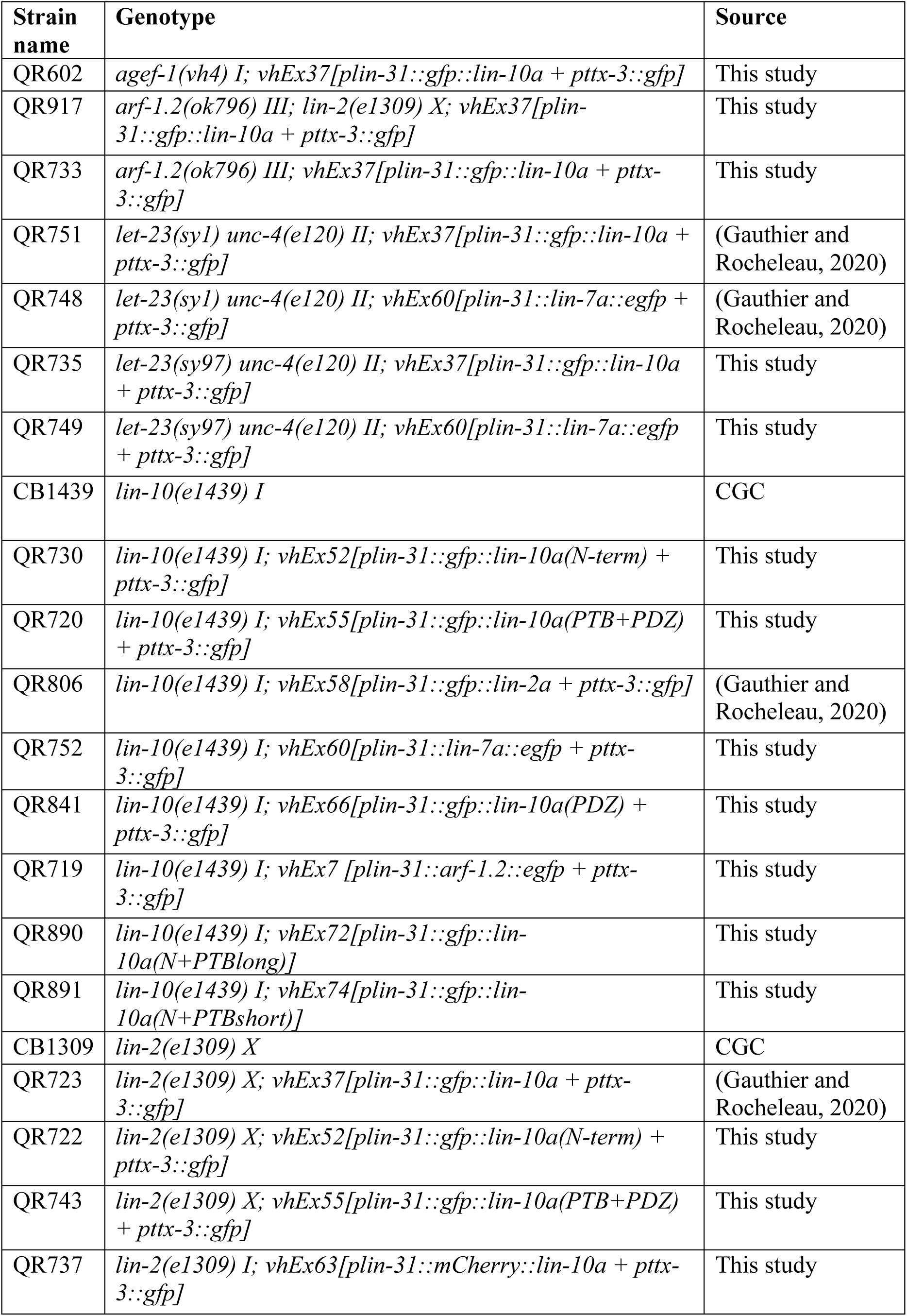
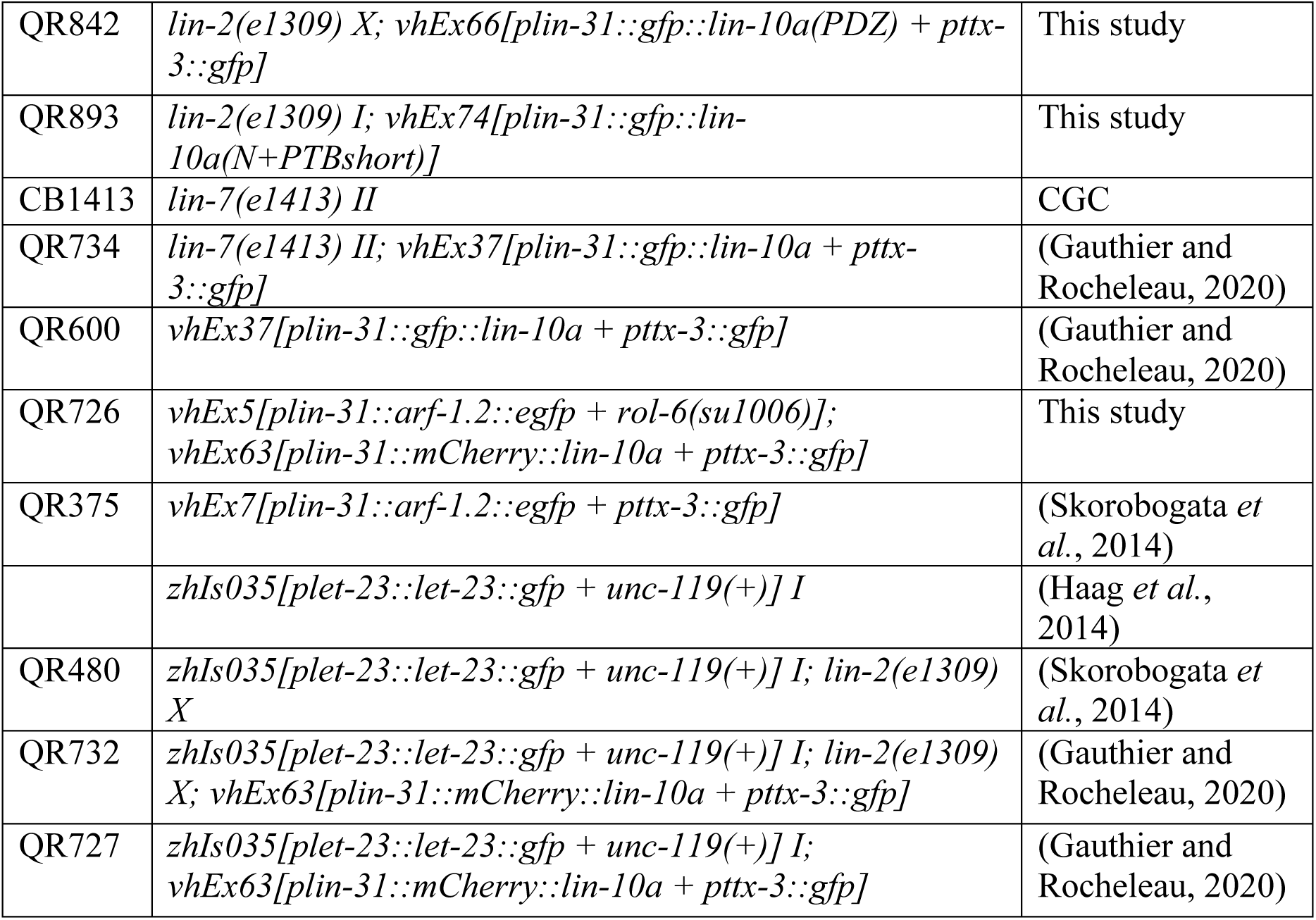
Strain list.

**Supplementary Table 2:**
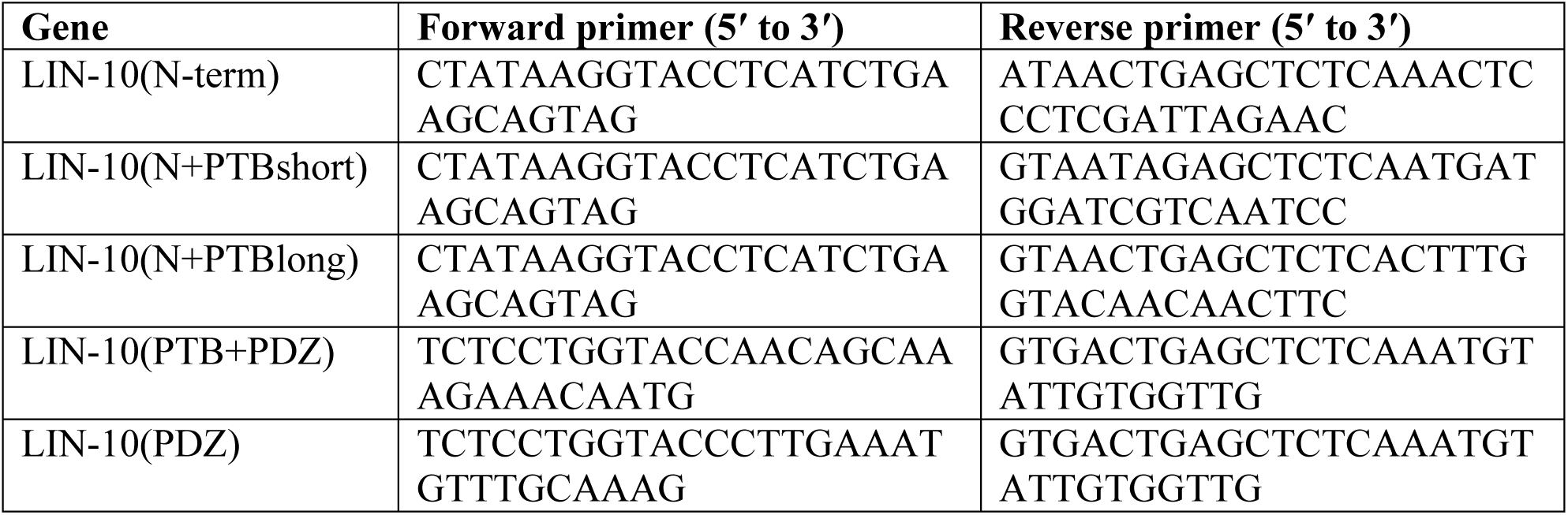
Primers for cloning.

